# Transpiration Responses to Potential Volatile Signals and Hydraulic Failure in Single Leaves of *Vitis Vinifera* (CV. Shiraz) and *Arabidopsis Thaliana* (Col 0) Utilising Sensitive Liquid Flow and Simultaneous Gas Exchange

**DOI:** 10.1101/2023.01.24.525440

**Authors:** Suzanne L. Balacey, Dimitra Liacopoulos Capone, Wendy Sullivan, Stephen D. Tyerman

## Abstract

Volatile organic compounds (VOCs) may communicate stress between plants. However little appears to be documented on how such VOCs affect transpiration. Changes in transpiration in response to some VOCs was examined by measurement of flow (*Q*) at high resolution into detached leaves of *Vitis vinifera* (cv. Shiraz) and Arabidopsis (Col 0). Sensors recorded arrival and decay of volatiles at the leaf lamina. Moderate xylem tensions were developed in *V. vinifera* leaves by incorporating a hydraulic resistance in the flow pathway. Simultaneous recording of leaf gas exchange (Assimilation, *A*, and Transpiration, *E*) for both *V. vinifera* and Arabidopsis revealed that for Arabidopsis *Q* was stochastically restricted by the gas exchange cuvette but not *E* in the short term. Depending on the applied supply pressure cavitation could be controlled in *V. vinifera* evident from reduced *Q*, and leaf wilting. Stomatal closure occurred upon cavitation after a transitory increase in *E* and *A*, and after wilting began and was reversed by re-pressurization. VOCs were emitted from leaves corresponding to changes in flow rate, and light to dark transitions but not to cavitation. Volatile methanol but not ethanol or methyl salicylate induced a localised dose-dependent reversible stomatal closure in both *V. vinifera* and Arabidopsis.

## 1. INTRODUCTION

By regulating the aperture of stomata, plants adjust transpiration and the intake of CO_2_ for photosynthesis when responding to various factors (Hetherington & Woodland, 2003) such as water availability (Buckley, 2019; Hernandez-Santana, Rodriguez-Dominguez, Fernández, & Diaz-Espejo, 2016; Tombesi et al., 2015), temperature (Urban, Ingwers, McGuire, & Teskey, 2017; von Caemmerer & Evans, 2015), light (Inoue & Kinoshita, 2017; Shimazaki, Doi, Assmann, & Kinoshita, 2007), CO_2_ concentration (Engineer et al., 2016; Israelsson et al., 2006), biotic stress (Melotto, Underwood, Koczan, Nomura, & He, 2006) and volatile organic compounds (VOCs) in the atmosphere (Jiang, Ye, Rasulov, & Niinemets, 2020; Niinemets & Reichstein, 2003). They are also the pathway for the release of some VOCs including simple small molecules such as methanol (Hüve et al., 2007). Hence, studying stomatal movement has been scrutinised in order to understand how plants respond to their environment and to potentially propagate VOC signals (Niinemets & Reichstein, 2003). There is less information available on the real time responses of stomata to potential VOC signals (López-Gresa et al., 2018).

Stomatal regulation can be monitored with various methods such as using stirred or unstirred chambers attached to the leaf; e.g. gas-exchange portable photosynthetic systems (Ceciliato et al., 2019; Jiang et al., 2020; Rasulov, Talts, & Niinemets, 2019) and porometers (Toro, Flexas, & Escalona, 2019), or with leaf isolated epidermal peels (Mott, Berg, Hunt, & Peak, 2014). These methods have been extensively used to improve the understanding of how stomata are regulated but most of these require direct contact with the leaf and isolation of the atmosphere over the measurement area of the leaf, making it difficult to study the impacts of VOCs.

Plants may use certain VOCs to communicate abiotic stress (Balacey, 2021; Midzi et al., 2022) and sophisticated systems have been developed to detect the *volatilome* from plants in real time (Jud, Winkler, Niederbacher, Niederbacher, & Schnitzler, 2018; Tholl, Hossain, Weinhold, Rose, & Wei, 2021). Soil born VOCs may also participate in interplant (root to root) signalling to stomata (Falik et al., 2012; Falik, Mordoch, Quansah, Fait, & Novoplansky, 2011). Our study developed methods to examine the leaf level transpirational responses from measurement of the flow into a detached leaf through the petiole in order to investigate responses to VOCs. Comparisons were made with simultaneously applied conventional leaf gas exchange. The methods were applied to both *Vitis vinifera* and *Arabidopsis thaliana* to validate reproducible results of continuous and homogeneous water flow rates (*Q*) into single leaves over time. Different artificial sap solutions feeding the leaf were trialled to optimise *Q* measurements. Comparisons with conventional gas-exchange (infra-red gas analyser (IRGA) systems) were conducted simultaneously to assess the water entering and leaving the leaf and to determine if the effects of VOCs or air movement were local or systemic though the leaf. Incorporation of a realistic hydraulic resistance in the flow pathway was examined for the development of negative leaf xylem pressures in *V. vinifera*. Light to dark transitions and hydraulic failure were examined as a reproducible way to test the responsiveness of stomata and to compare with literature data (Elhaddad, Hunt, Sloan, & Gray, 2014; Jardine et al., 2012).

Possible plant VOCs that may act as signals for plants were tested for their effect on transpiration. For instance, López-Gresa et al. (2018) showed that four hexenyl esters ((*Z*)-3-hexenyl acetate, (*Z*)-3-hexenyl propionate, (*Z*)-3-hexenyl butyrate, and (*Z*)-3-hexenyl isobutyrate) were responsible for the closure of tomato stomata in response to a pathogen attack. Other common plant-emitted VOCs were tested such as ethanol (Jud et al., 2016) and methanol (Folkers et al., 2008; Jud et al., 2018); the latter causing reversible dose-dependent stomatal closure. In addition, the technique was improved by monitoring VOCs in real time with a relatively cheap but non-specific sensor during treatments and/or those emitted by the leaf.

## 2. MATERIALS AND METHODS

### 2.1 Plant and environmental conditions

*Arabidopsis thaliana* Col 0 were potted and grown in a small growth cabinet with artificial light (PAR 100-150 μmol m^−2^ s^−1^) for 5 weeks with short-day environmental conditions (10 h light, 21°C / 14 h dark, 17°C; humidity 60 %). *Vitis vinifera* L. cv. Shiraz (Syrah) vines were potted and grown in a glasshouse under natural light until reaching 1-2 shoots with approximately 10 leaves per shoot. The environmental conditions were temperature 25°C day and 17°C night, humidity 40 %. Shiraz leaves were also taken from the field during spring growth as indicated.

### 2.2 Flow meter parameters

Flow rate (*Q*) into single leaves was monitored using a modified XYL’EM embolism meter (Instrutec, France) with added high precision liquid flow meters (LIQUI-FLOW, Bronkhorst High-Tech B.V., Netherlands) (Cochard, 2002; Cochard, Bodet, Améglio, & Cruiziat, 2000; Cochard, Coll, Le Roux, & Améglio, 2002), at different sensitivities depending on the species being measured (*V. vinifera*, 5 g h^−1^; Arabidopsis, 0.5 g h^−1^ maximum flow rate) (Figure 1). Through the dedicated software ‘XYL_WIN’, flow rate (g h^−1^), temperature and pressure were recorded at selected time intervals, usually every 10 s. The instrument was filled with a solution of purified water (Milli-Q Plus; Merck Millipore, Billerica, MA, USA) which was degassed (1.0 × 5.5 Mini Module ™; Membrana GmbH, Germany) to avoid formation of air bubbles. High pressure tubing of sufficient volume (depending on flow rate and total time of the experiment) was incorporated in the flow pathway containing various artificial saps. PEEK capillary tubing (length 3 m, internal diameter 0.2 mm; hydraulic conductance = 7.65 × 10^−6^ kg s^−1^ MPa^−1^) was included in series in some of the experiments with *V. vinifera* to induce more negative xylem pressures (Figure 1). This was similar to the average hydraulic conductance of the remainder of the petiole in series (11.1 × 10^−6^ kg s^−1^ MPa^−1^, SD = 7.2 × 10^−6^ kg s^−1^ MPa^−1^, n=24) of average length 49 mm (SD= 10.2, n=24). The leaf petiole of *V. vinifera* was sealed at the end of the PEEK tubing with a silicone tubing washer (Mastreflex L/S, Precision Pump Tubing, C-FLEX, L/S, Cole-Parmer, USA) inside a gas tight fitting. In these experiments variable pressures were applied at the up-stream end (at the instrument) to control the xylem tension at the leaf petiole.

**Figure 1.**
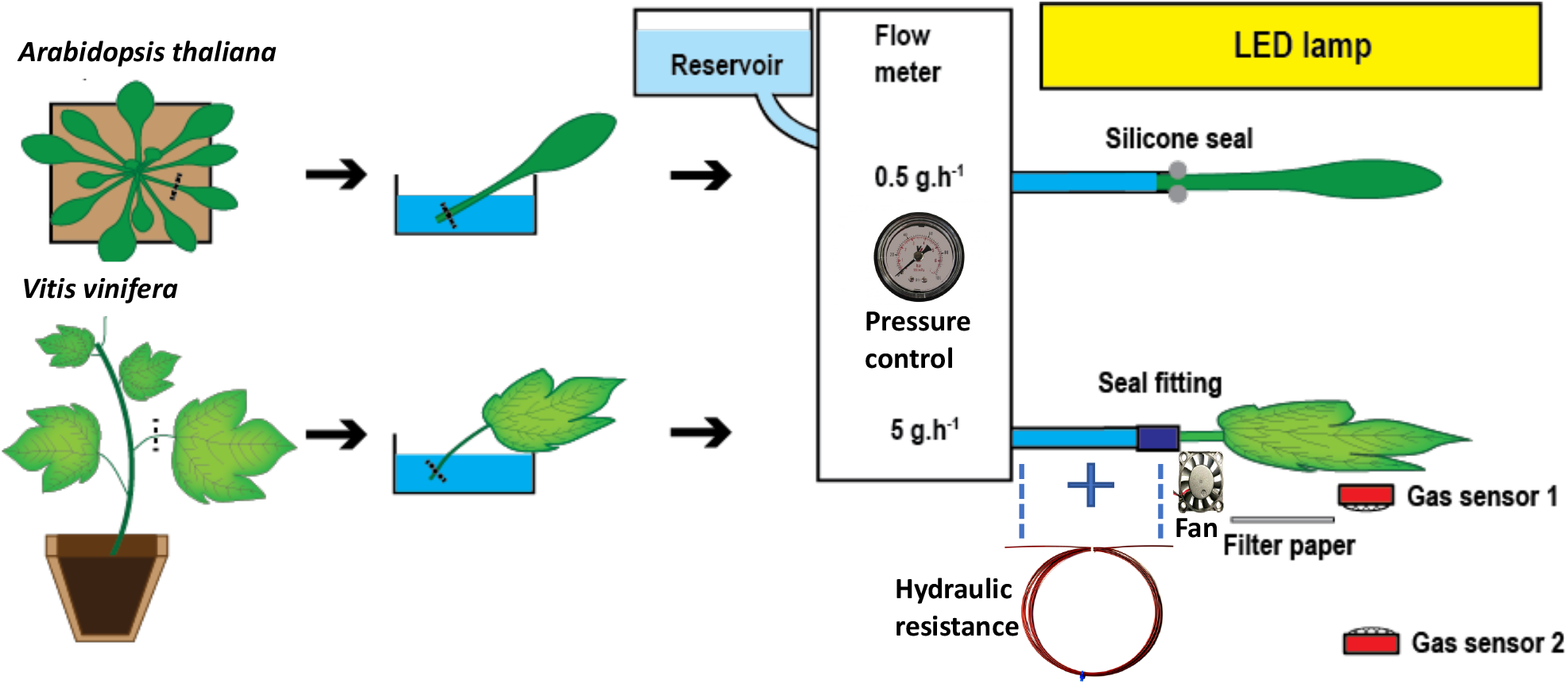
Depiction of the method used to perform whole-leaf real-time-resolved analyses of stomatal responses to volatile compounds, xylem tension and airflow in *Arabidopsis thaliana* and *Vitis vinifera*. Leaves were severed in the artificial sap and connected to a flow meter for continuous measurements of water flow (*Q*, mmol m^−2^ s^−1^) under a LED lamp (PAR 150 – 1300 μmol m^−2^ s^−1^ at leaf level depending on experiment). A hydraulic resistance was incorporated in experiments with *V. vinifera* to induce tensions in the leaf xylem depending on the supply pressure controlled by the instrument. VOCs were added underneath the leaf on a filter paper and monitored with gas sensors to follow their effect on stomatal responses. A fan was also trialed to examine whether it could affect flow into the leaf by reducing the boundary layer resistance.

Artificial saps were composed of purified degassed water with either: 1) 10 mM KCl, 2) 10 mM KNO_3_, 1 mM MES (2-(N-Morpholino)-ethanesulfonic acid; pH 5.5), or 3) 3 mM KNO_3_, 0.1 mM MgSO_4_, 1 mM CaCl_2_, 1 mM KH_2_PO_4_, 1 mM K_2_HPO_4_, pH 6 with HCl (Wilkinson and Davies artificial sap, WD-AS)) (Wilkinson & Davies, 1997). These were filtered through a 0.2 μm syringe filtration unit (Filtropur S 0.2, Sarstedt, Germany).

### 2.3 *Vitis vinifera* measurements

Fully expanded undamaged mature leaves were randomly selected from *V. vinifera* at similar growth stages (between nodes 3-6). They were severed from the shoots by cutting with a pair of sharp scissors (about 2-3 mm distance to the stem junction) and immediately immersed in a petri dish filled with the artificial sap solution. The petiole was recut with a razor blade to avoid risk of embolism and wiithin a minute, the petiole was tightly sealed with plastic fittings attached to the tubing of the flow meter. Light was provided by a dedicated photosynthetic LED light (Mars Reflector 48, Mars Hydro, USA) placed between 15 and 40 cm over the leaf providing between approximately 150 to 1300 μmol m^−2^ s^−1^ PAR at leaf level respectively depending on the particular experiment. Experiments began between 10 am and 12 noon and were usually completed by 5 pm. Once the measurements were complete, the projected leaf area was calculated from scans with ImageJ (FIJI) to determine the water flow rates (*Q*) in mmol m^− 2^ s^−1^. In some experiments where the extra hydraulic resistance was incorporated in the flow pathway, the remaining hydraulic conductance in the petiole was measured. This allowed a total tube-petiole hydraulic conductance (*K*_t_, mmol s^−1^ MPa^−1^) to be obtained and this was used to calculate a xylem pressure (*Px*) at the leaf entry point depending on measured flow rate (*Q*):

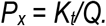

### 2.4 Time lapse photography

To record leaf wilting a Sony (HDR-AS30V) camera was set to take images every 60s. ImageJ (FIJI) was used to measure a leaf tip position over time using the image stack and point tool. This is plotted as a relative change in position, with zero being fully wilted and 1 being fully turgid.

### 2.5 *Arabidopsis thaliana* measurements

A similar protocol was utilised for Arabidopsis leaves, but with slight modifications. The petiole was directly cut with a fresh razor blade, and quickly immersed in a petri dish with artificial sap. The second cut was performed following the recommendations described in (Ceciliato et al., 2019), i.e. by gently moving the razor blade back and forth and not pressing the blade against the petiole, potentially damaging the xylem conduits. The tubing was sealed with silicone paste (Xantopren L blue, Heraeus, Germany) due the petiole not being circular in cross section and thus it could not sustain a pressure seal.

### 2.6 Simultaneous measurements with a gas exchange instrument

A LCpro-SD portable photosynthesis system (ADC BioScientific Ltd., UK) was added to enclose a portion of the leaf for *V. vinifera* and the whole leaf for Arabidopsis (sealed at the petiole), supplied with ambient CO_2_ concentration, temperature and humidity, at an air flow of 300 mL min^−1^ in order to measure transpiration (*E*) and assimilation (*A*) rates. A clear top LCpro-SD head was used so light was provided by the LED lamp over the leaf. Data acquisition was programmed with automatically timed-logging.

### 2.7 Volatile application

A range of volatiles were tested to examine their impact on water flow rates whilst the leaf was connected to the flow meter and/or with simultaneous gas exchange. A piece of cellulose filter paper (Whatman plc, UK) was placed approximately 2-3 cm underneath the leaf and the volatile chemical of interest was pipetted with different concentrations and volumes as described in Table 1. The concentrations of the esters were chosen according to (López-Gresa et al., 2018).

**Table 1.** Volatile compounds selected, concentrations and volumes added on the filter papers.

| Volatile compounds | Concentration | Volumes added to the filter paper (µL) |
| --- | --- | --- |
| ethanol (EtOH) | 100 % | 150, 200, 1000, |
| methanol (MeOH) | 100 % | 10, 20, 50, 100, 200 500 |
| methyl salicylate | 100% | 100 |
| (Z)-3-hexenyl butyrate | 5 µM in ethanol (100 %) | 200 |
| (Z)-3-hexenyl propionate | 5 µM in ethanol (100 %) | 200 |
| (Z)-3-hexenyl isobutyrate | 5 µM in ethanol (100 %) | 200 |
| (Z)-3-hexenyl acetate | 5 µM in ethanol (100 %) | 200 |

### 2.8 Sensors and parameters

A temperature/humidity sensor (DHT22, Aosong (Guangzhou) Electronics Co.,Ltd, China) and two VOC sensors (Grove-gas sensor MQ3, Seeed studio) were added to the system (Ionescu & Vancu, 1996). One gas sensor (VOC1) was placed either underneath the leaf (approx. 10 mm) or in parallel with the leaf surface to the side of the leaf blade (due to the sensor emitting heat) while another was placed 15 - 30 cm below to distinguish between the origins of the volatiles (e.g. emitted by the leaf or coming from the surrounding environment) (Figure 1). The MQ3 sensor is composed of micro aluminium oxide (Al_2_0_3_) ceramic tube, tin dioxide (SnO_2_) sensitive layer, measuring electrode and heater, fixed into a small chamber and is highly sensitive to organic solvent vapours such as ethanol and methanol but is not specific. The VOC sensors were connected to an analog digital converter (10 bit) inputs of an Arduino UNO microcontroller to record measurements to an SD card on a Adafruit Data Logging shield (Coding in Supplementary Material S2). The DHT22 sensor was connected to digital input pins and recorded simultaneously with the gas sensors. Time shown on x-axes are Australian Central Standard Time (AusCST) with 12:00 approximately corresponding to solar noon in the local environment. All instrument time was synchronised to the nearest second using a smart Phone timer App.

### 2.9 Gas sensor calibration

Both sensors were calibrated against methanol solutions in pure distilled water (50 mL) in a closed (stirred air) container at a constant temperature and humidity. Humidity does effect the calibration but could only be performed at about 90% RH. During experiments humidity will be less than this, thus this could only contribute a small effect. Henry’s law was used to calculate methanol vapour concentration for a given temperature. The sensors give a logarithmic voltage output (Voltage = A * (ppmv) ^B^) as a function of gas concentration (where A and B are coefficients determined from regression), and this resulted in calibration curves with an R^2^ > 0.99 could be obtained (Supplementary Figure S1). Previous experiments have shown high predictability using this calibration against unknown solutions of ethanol subsequently measured enzymatically.

### 2.10 Fan application to *V. vinifera* leaves

A fan (80 mm diameter; Panaflo, model FBA08A24H, air volume 1.87 × 10^−2^ m^3^ s^−1^, Panasonic, Japan) was added to the system to test the effect of altering the boundary layer resistance. The airflow was directed perpendicular and on to the abaxial leaf surface. Alternatively a small (30 mm diameter, 5 volt) fan (Sirocco YX201) was used to generate air flow (approx. 1.45 m^−3^ s^−1^) directed parallel to and across the abaxial surface of the leaf.

## 3. RESULTS

### 3.1 Water flow into single leaves measured with the flow meter

After connecting a leaf to the flow meter, it took approximately 30 - 120 min for *V. vinifera* leaves and 20 - 30 min for Arabidopsis leaves to reach a plateau and stabilise with the flow rates in the light averaging 1.94 ± 0.6 mmol m^−2^ s^−1^ (mean ± SD, n=47) for *V. vinifera* (Figure 2) and 1.05 ± 0.29 mmol m^−2^ s^−1^ (mean ± SD, n=9) for Arabidopsis leaves. Maximum flow rates for *V. vinifera* could reach about 5 mmol m^−2^ s^−1^ at higher light intensities. For *V. vinifera* leaves perfused with WD-AS measurements were generally stable over time (Figure 2a), while for Arabidopsis flow reached a maximum then decreased to a relatively stable level (Figure 2b). Three artificial saps were tested in the flow meter system. High and similar initial flow rates were obtained with all sap solutions (Figure 2) and either the MES/KNO_3_ artificial sap solution or the WD-AS were used for further experiments as indicated. The WD-AS solution gave more steady flow rates over time.

**Figure 2.**
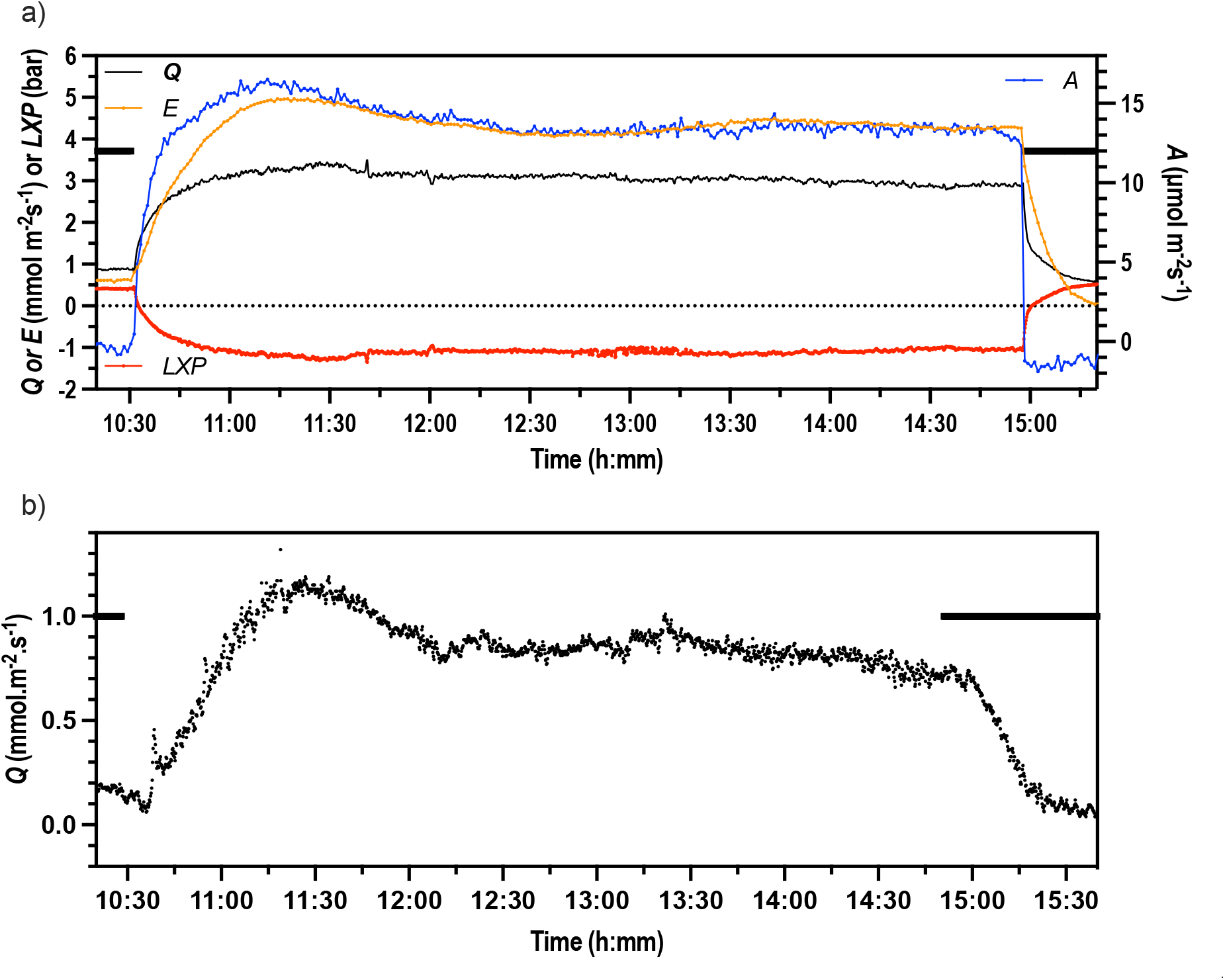
Flow rate (*Q*) measured in a) *V. vinifera* (cv. Shiraz) leaf (field collected) and b) Arabidopsis leaf connected to a liquid flow meter demonstrating dark-light-dark transitions (solid black line = dark) and relatively stable flow rates. a) For *V. vinifera* in this instance simultaneous gas exchange was measured (*E* orange points, *A* blue points) and a hydraulic resistance was incorporated to generate tension within the leaf xylem (Leaf Xylem Pressure, *LXP*, red points) (PAR = 1170 μmol m^−2^ s^−1^). B) Arabidopsis leaf using the more sensitive flow meter (PAR = 100 μmol m^−2^ s^−1^). It was not possible to perform simultaneous gas exchange for reasons given below. For both the WD-AS was used (pH 6 for *V. vinifera*, pH 4.5 for Arabidopsis).

### 3.2 Comparison of flow rates (*Q*) with transpiration rates (*E*) utilising a gas-exchange system

As the flow meter is measuring the water entering the leaf via the xylem, the water that is transpired through stomata was measured by connecting an Infrared Gas Analyser (IRGA) photosynthetic system (LCpro-SD) for comparison. In order to assess the effect of clamping the IRGA cuvette to a leaf, *Q* was initially measured until steady before the cuvette of the IRGA was connected. For Arabidopsis, as soon as the IRGA cuvette was clamped on the leaf, the *Q* measured with the flow meter was disrupted and displayed fast oscillations around approximately 0 mmol m^−2^ s^−1^, but returned to initial values once the cuvette was removed (Figure 3a). The IRGA recorded similar *E* as *Q* from the flow meter measured prior to the attachment of the head, starting at 0.5 mmol m^−2^ s^−1^. The dark transition induced a decrease in *E* measured with the IRGA and an increase was observed once the light was turned on again. Other examples are shown in Supplementary Figure 2.

**Figure 3.**
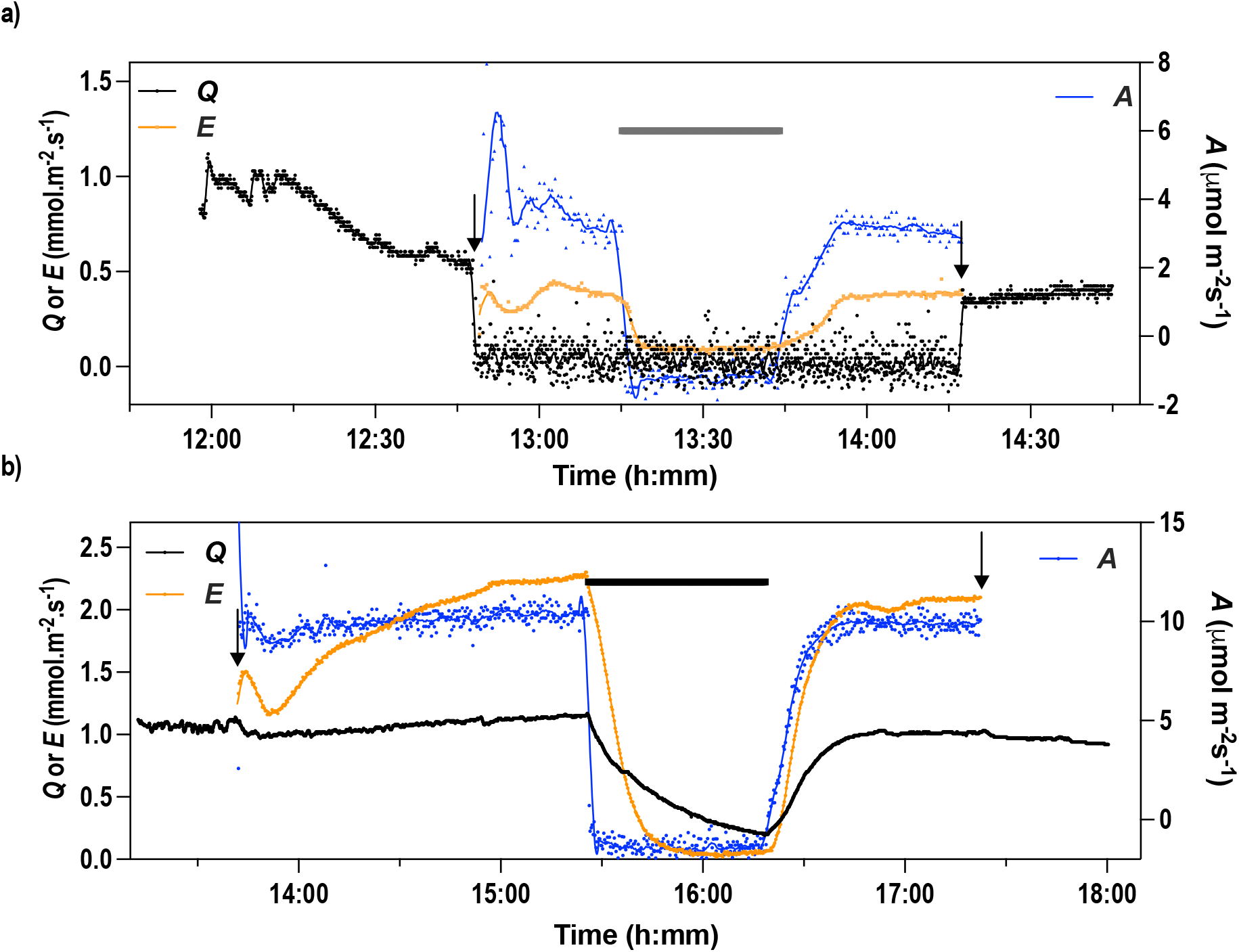
Effect of connecting the gas exchange cuvette while simultaneously measuring flow (*Q*, black points) into the leaf. Measurements of *Q*, transpiration (*E*, orange points) and assimilation (*A*, blue points) for a) *Arabidopsis thaliana* Col 0 and b) *V. vinifera* (cv. Shiraz, glasshouse grown). The arrows indicate when the gas analyser cuvette was attached and removed from the leaf. Dark transitions are indicated (black horizontal line, approx. PAR 10 μmol m^−2^ s^−1^). Sap solution = 10 mM KNO_3_ pH 5.5, PAR = 260 (a) and (b) 290 μmol m^−2^ s^−1^. For both species, data shown are from one representative experiment out of 3 repetitions with similar results.

For *V. vinifera*, clamping the cuvette on the leaf disrupted *Q* measured with the flow meter but not as much as for Arabidopsis (Figure 3b). *E* measured with the IRGA reached a twofold higher rate than *Q* after 1 h in this example, but this difference varied between experiments either above or below *Q*. Note that the fraction of leaf area in the IRGA (6.25 cm^2^) is relatively small compared to the total mean leaf area of 159 cm^2^ (+/− 35 SD, n=23) for which *Q* is measured simultaneously. The area of the leaf under the IRGA cuvette seals was subtracted in calculating the area for *Q*. The dark transition induced a decrease of *E* and *Q*, but with different rates.

Oscillations were sometimes observed in *Q* after dark-light transitions or after attaching the leaf initially under the light and these were reflected also in *E* and *A* when gas exchange was recorded simultaneously. Oscillations in *Q* and *E* had about the same frequency but were not synchronous, as peaks in *E* and *A* preceded peaks in *Q* (Figure 4). Other examples are shown in Supplementary Figure 3.

**Figure 4.**
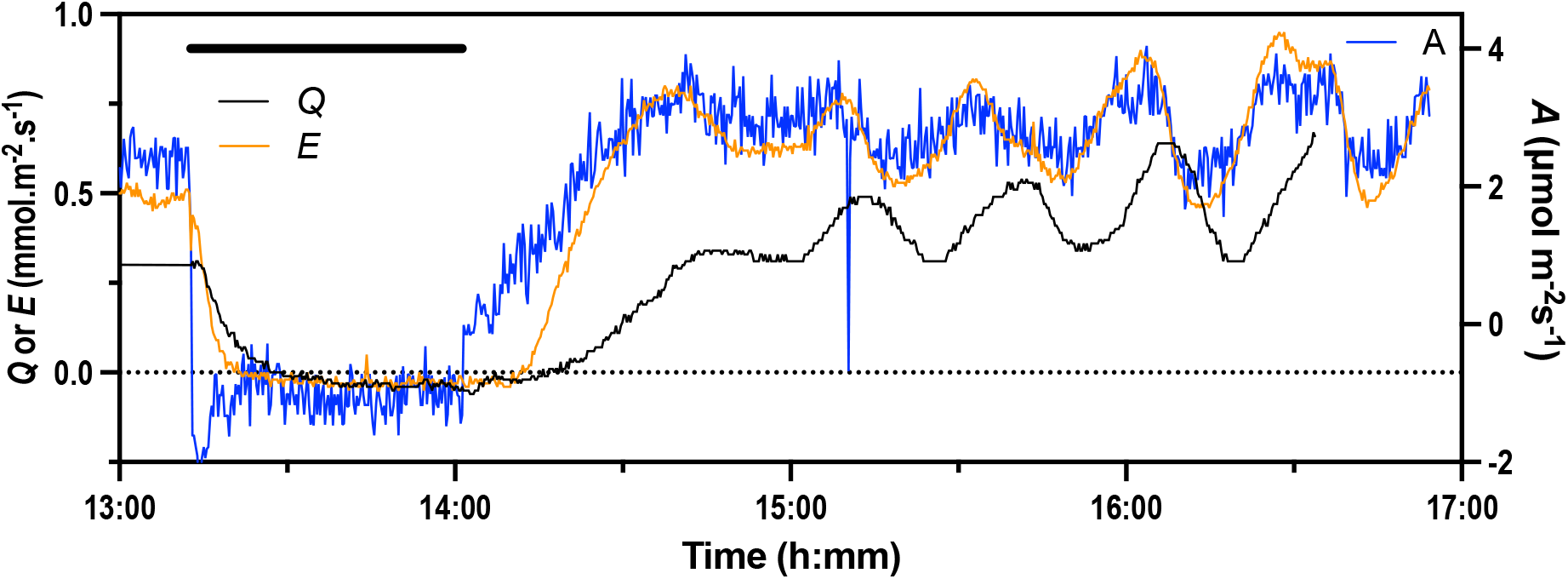
Oscillations observed in flow (*Q*) and gas exchange (*E* orange points, *A* blue points) in *V. vinifera* (cv. Shiraz) (PAR = 720 μmol m^−2^ s^−1^). In this case the leaf was already attached to the flow meter and IRGA for 24 hours and a dark transition was applied (black horizontal line, approx. PAR 10 μmol m^−2^ s^−1^). Sap solution = 10 mM KNO_3_ pH 5.5.

### 3.3 Effects of dark-light transitions on flow rates

To examine the stomatal responses of the leaf, dark-light-dark transitions were conducted by turning the LED lamp positioned above the leaf on and off. For *V. vinifera* a dark-light transient caused the flow to initially dip slightly then rapidly increase prior to a more gradual increase to a relatively constant flow (e.g. Figure 2a, Figures, 4–9). The example shown in Figure 2a had gas exchange simultaneously measured and had the flow resistance incorporated (see below). For both *V. vinifera* and Arabidopsis, the response in flow after the light was switched on or off was almost instant. For light-off Arabidopsis *Q* reached approximately 0 with a half-life of 1020 ± 777 s (n=3). For *V. vinifera*, the half-life was 309 ± 56 s (n=3). Dark-light-dark transitions were used to assess the responsiveness of the stomata while being connected to the flow meter in further experiments with VOCs.

### 3.4 Leaf xylem tension and hydraulic failure in *V. vinifera* leaves

By incorporating a hydraulic resistance in the flow path and measuring petiole hydraulic resistance at the end of the experiment it was possible to deduce the leaf xylem pressure (*LXP)* at least proximal to the attachment of the petiole to the leaf (e.g. Figure 2a). The feed pressure in the instrument could be varied to increase or decrease this tension and from prior experiments it could be approximately estimated during an experiment. An example is shown in Figure 5 where the leaf was also monitored with time lapse photography to record wilting. In this case the feed pressure was gradually reduced (spikes in *Q* in Figure 5) until just after the last reduction when *LXP* reached about −1.5 bar and *Q* suddenly decreased (first arrow in Figure 5). Almost simultaneously the leaf started to wilt. This corresponded to a transitory increase in *E* and *A* before a relatively rapid reduction. This transitory increase was always observed and could be up to 1.5 × the initial *E* and *A* values. Re-pressurization resulted in the leaf regaining turgor and *E* and *A* partially recovering; note that the calculated *LXP* remained slightly negative during this process. Turning the light out in this case led to further recovery of leaf turgor as indicated by a further rise in the leaf tip position (*LTP*). Hydraulic failure could also be induced by increasing airflow across the leaf (Supplementary Figure 4).

**Figure 5.**
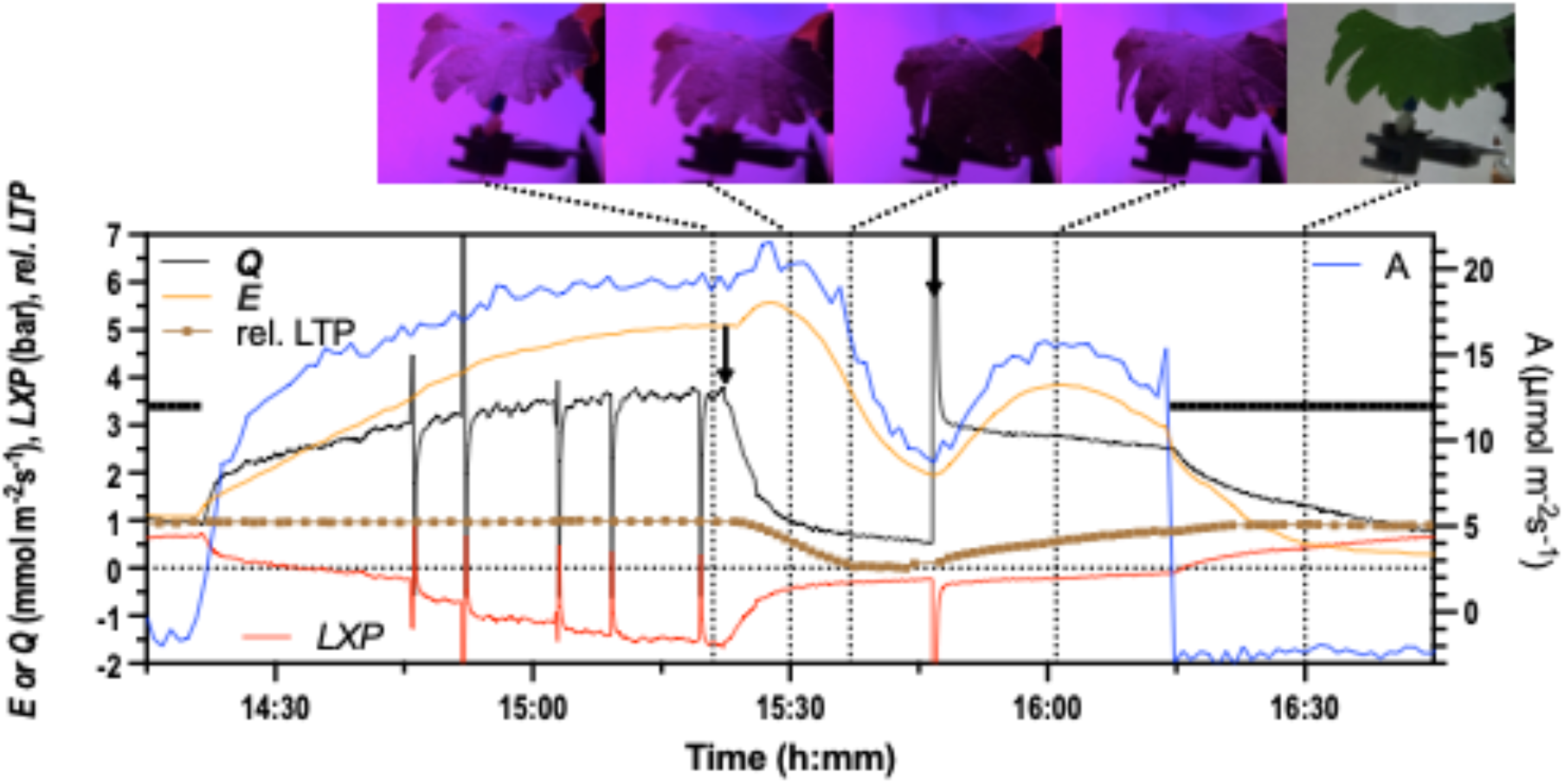
Effect of hydraulic failure on flow, gas exchange and leaf wilting in *V. vinifera* cv. Shiraz. In this case the hydraulic resistance was incorporated in the flow path to the leaf resulting in significant negative leaf xylem pressures (*LXP*). Progressive lowering of the feed pressure resulted in increasing tensions to the point where flow collapsed (first arrow). This corresponded to the start of leaf wilting indicated as relative leaf tip position (rel. *LTP* brown points and line) and corresponding images of the leaf above. At the second arrow the feed pressure was increased restoring flow and resulting in *E*, *A* and leaf turgor recovering. Sap solution = WD-AS pH 6.0, PAR = 770 μmol m^−2^ s^−1^.

During wilting and recovery there was no evidence for a detected VOC signal arising from the leaf (n=4), though on other occasions as outlined below changes in flow and light-dark transition did result in VOC emission.

### 3.5 Effect of air flow on water flow rates

To further examine the response of *Q* to leaf transpiration a fan was installed under the leaf to test if increased air movement by decreasing the leaf boundary layer resistance would have the anticipated effect on *Q*, i.e. greater air flow would reduce the boundary layer resistance, which would increase Q and *vice versa*. The fan was turned on after obtaining a steady *Q*. This caused a rapid increase in flow followed by a slower decay back toward the initial rate sometimes over shooting (e.g. Figure 6, Supplementary Figure 4). This response was observed several times with variable decay rates, and it was decided that all remaining experiments would be conducted without the fan in order to record responses with the gas sensors to VOCs. Transpiration (*E*) measured with the IRGA (Figure 6a) decreased with increased air flow over the rest of the leaf and *vice versa*. This apparent systemic response in *E* could be attributed to a reduction in chamber and leaf temperature by about 3 °C while the fan was on.

**Figure 6.**
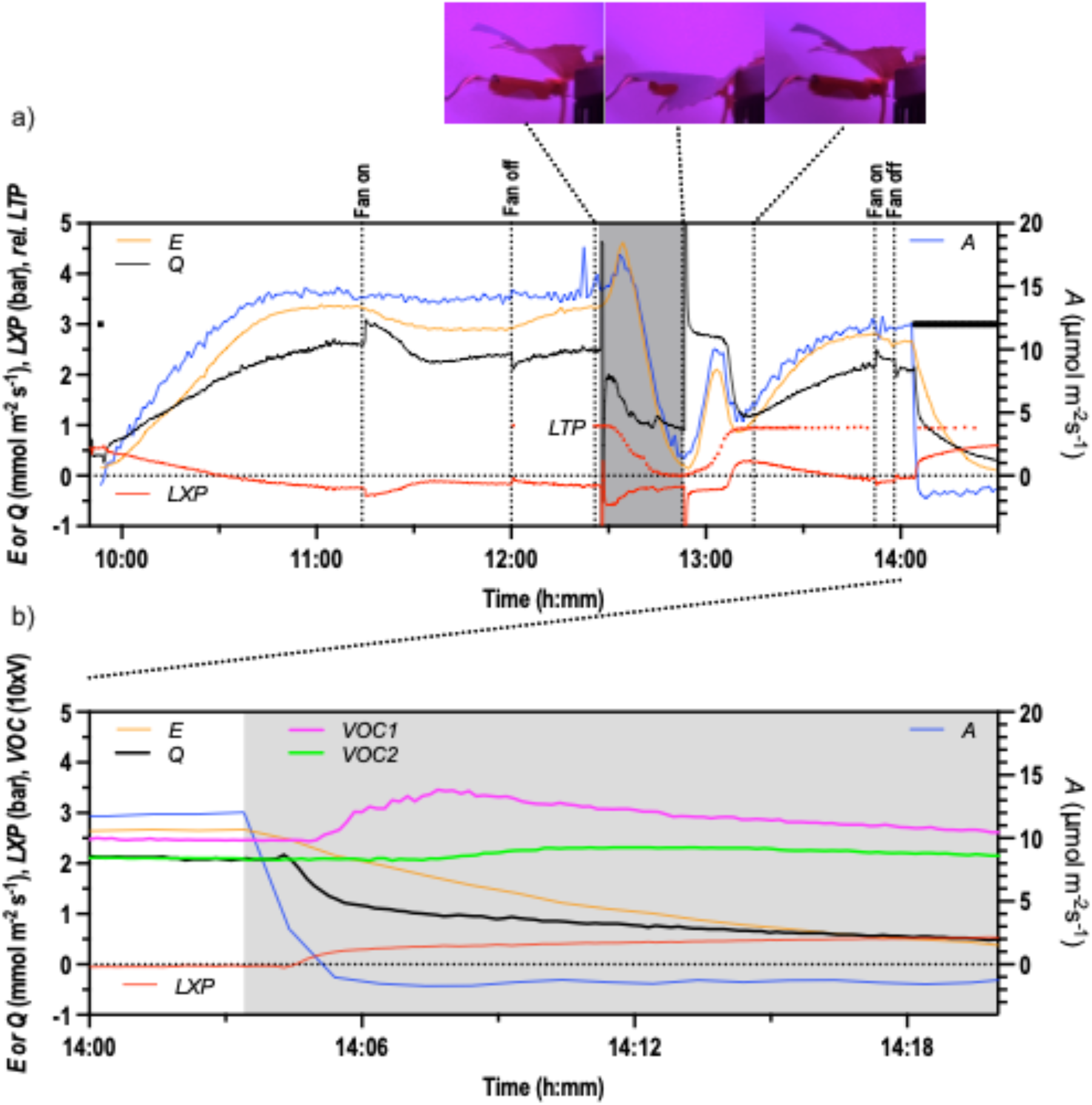
Effect of air flow over the leaf and hydraulic failure on flow, gas exchange and leaf wilting in *V. vinifera* cv. Shiraz (field grown) and example of volatile release upon stomatal closure with darkness. a) Turning the fan on resulted in an initial increase then gradual decrease (and overshoot) in flow. The reverse occurred when the fan was turned off. Note the response in *E* (orange points) but not *A* (blue points) and that the cuvette of the IRGA was isolated from the generated air flow from the fan. The hydraulic resistance incorporated in the flow path to the leaf resulted in negative leaf xylem pressures (*LXP* red points). The feed pressure was initially set at 0.7 bar and then dropped to 0.1 bar at the beginning of the shaded section. This induced hydraulic failure corresponded to the start of leaf wilting indicated as relative leaf tip position (rel. *LTP*) and corresponding sampled leaf images above. The feed pressure was re-established to 0.7 bar at the end of the shading. b) Detail of the light-dark transition where the VOC sensors picked up a signal emanating from the leaf as indicated by an earlier signal from the VOC1 sensor closest to the leaf (VOC1 magenta, VOC2 green). Sap solution = WD-AS pH 6.0, PAR = 920 μmol m^−2^ s^−1^.

### 3.6 Effect of VOCs on water flow rates

Ethanol was used as the solvent to dilute the hexenyl esters, and it was initially examined to determine its effect on *Q* for *Vitis vinifera* (Figure 7). A series of experiments showed inconsistent results where sometimes a small decrease of *Q* was induced and sometimes no effect (i.e. 2 out of 18 replications showed a response). The four hexenyl esters in ethanol ((*Z*)-3-hexenyl butyrate, (*Z*)-3-hexenyl isobutyrate, (*Z*)-3-hexenyl propionate, (*Z*)-hexenyl acetate) were also tested in duplicate at the same concentration as detailed in (López-Gresa et al., 2018) but they did not induce a reduction of *Q* (Figure 7a).

**Figure 7.**
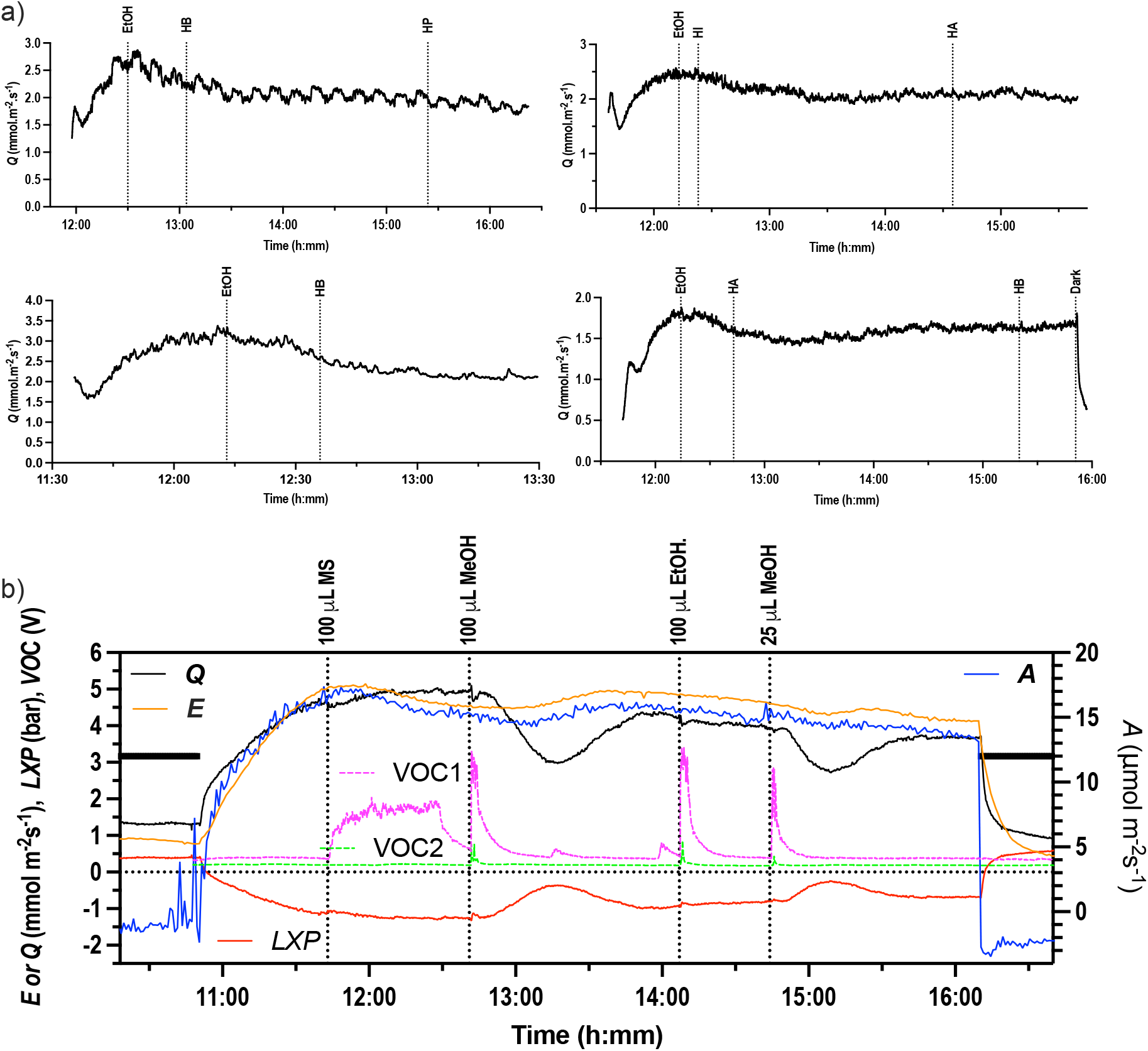
Effect of VOCs on flow rate (*Q*) into *V. vinifera* (cv. Shiraz) leaves. a) Four separate leaves glasshouse grown) under light: 200 μL each of ethanol (EtOH), hexenyl butyrate (HB, 5 μM (in ethanol)), hexenyl propionate (HP, 5 μM (in ethanol)), hexenyl isobutyrate (HI, 5 μM (in ethanol)), and hexenyl acetate (HA, 5 μM (in ethanol)) were added on a filter paper underneath the leaf. Sap solution = 10 mM KNO3 pH 5.5, PAR = 150 μmol m^−2^ s^−1^. b) An example of effect of VOCs (methyl salicylate, methanol and ethanol) where the VOC sensors were installed as shown in Figure 1 and gas exchange was measured simultaneously (field grown). Note that the gas exchange cuvette was isolated from the applied VOCs. In this case the hydraulic resistance was in the flow path and leaf xylem pressure (LXP) is shown. Sap solution = WD-AS pH 6.0, PAR = 1190 μmol m^−2^ s^−1^.

Methanol was also examined and different volumes of pure methanol (10 to 500 μL) were added onto a filter paper which was placed under *V. vinifera* leaves that were connected to the flow meter. The methanol treatments were monitored with gas sensors placed close to the leaf (parallel with lamina), and 15 - 30 cm under the leaf (gas sensor 1 and 2 in Figure 1). In every trial, the gas sensor 1 (VOC1 in Figures 6, 7, 9) sensed more methanol prior to gas sensor 2 (VOC2), and methanol stopped being detected after approximately 15 min application depending on the amount applied. For higher volumes of methanol there was an immediate small reduction then an increase in *Q* to the initial value that could be resolved (Figure 8a). This was inverse to the transient VOC concentration measured at the leaf level (Figure 8).

**Figure 8.**
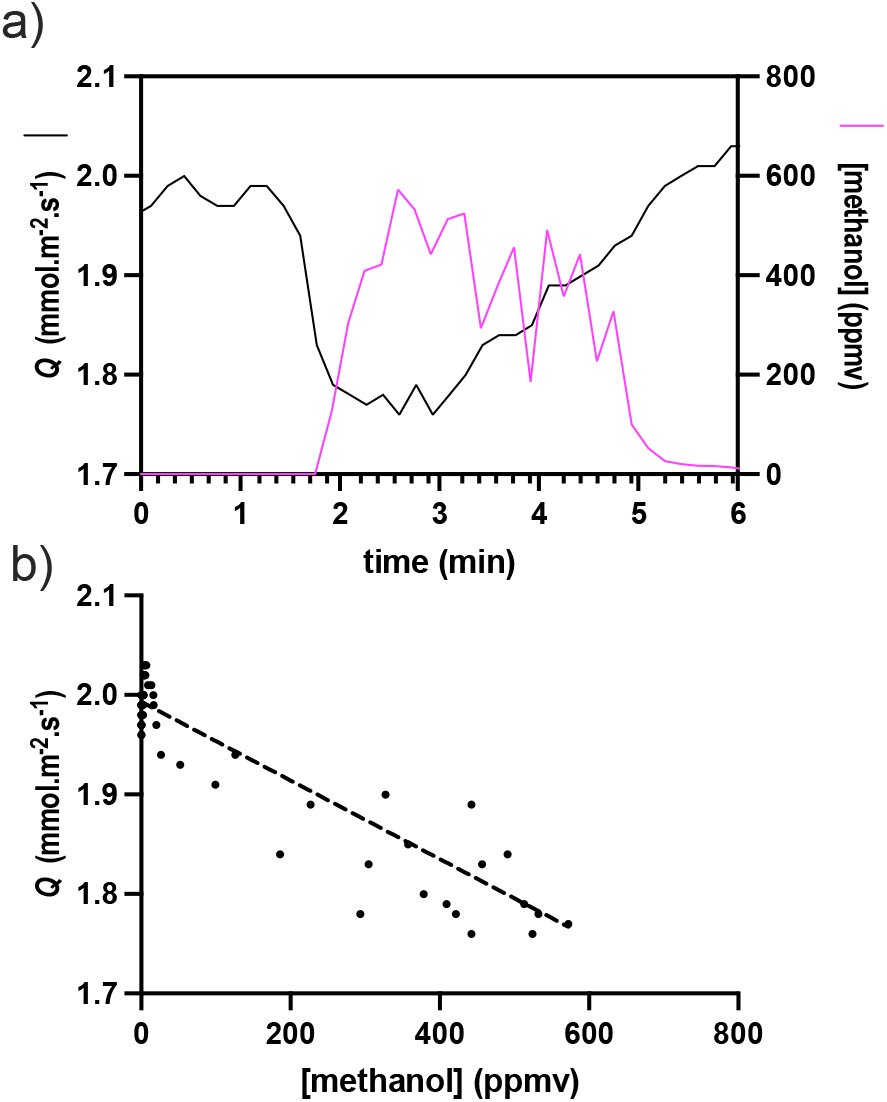
Initial transient in flow corresponding to methanol concentration estimated at the lamina from the VOC1 sensor where 200 μL of methanol was applied to the filter paper below a *V. vinifera* leaf. a) Transients in *Q* (black line) and methanol (magenta line) plotted against time. b) Regression of *Q* against methanol concentration illustrating negative correlation. R^2^ = 0.87, P<0.0001, Equation: *Q* = −4.243 × 10^−4^ × [methanol] + 1.993. The mean slope (n=3) was −3.211 × 10^−4^ mmol m^−2^ s^−1^ ppmv^−1^ (SD = 0.958 × 10^−4^)

**Figure 9.**
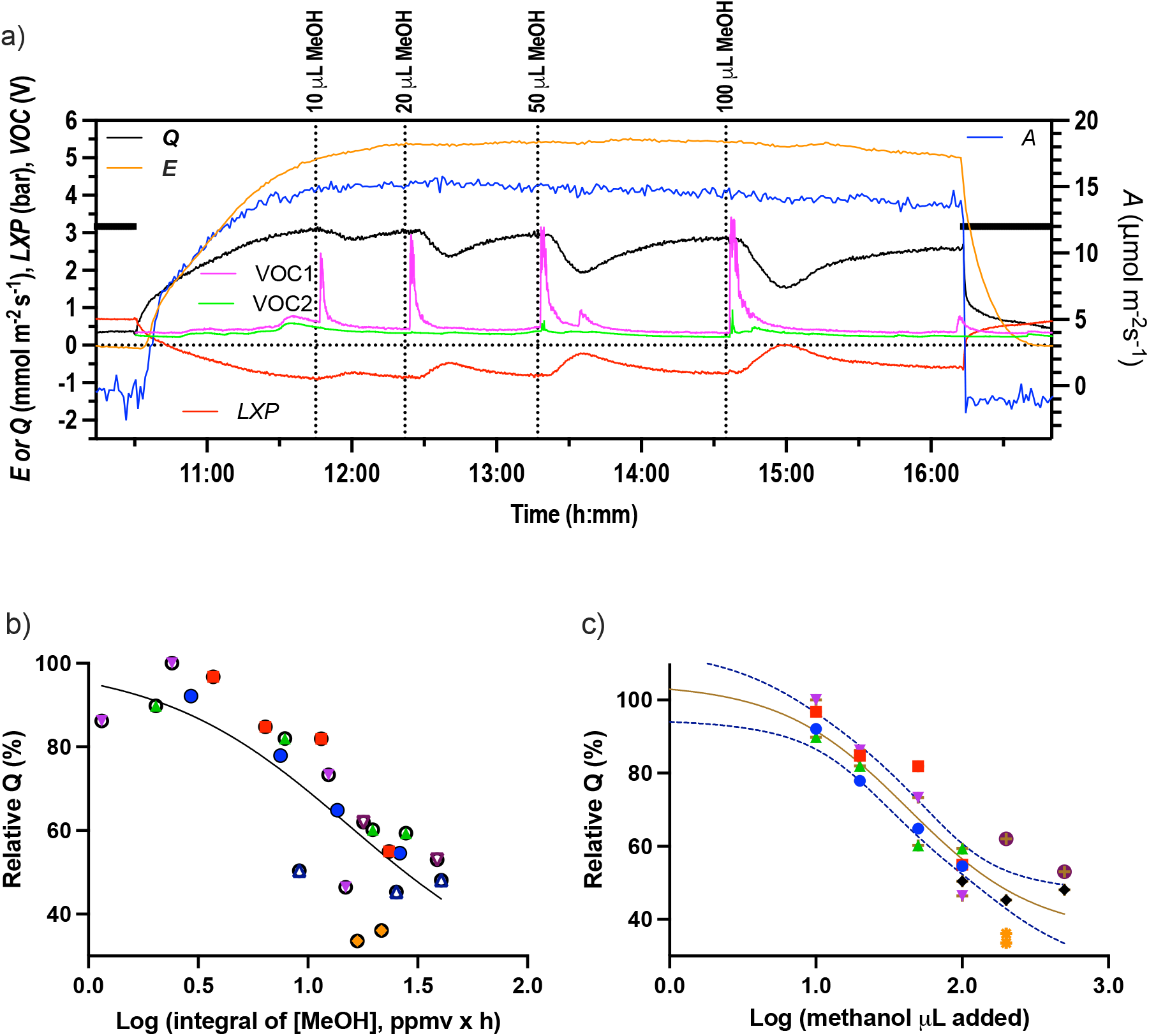
Methanol dose-response. a) Example of flow (*Q*) into a *V. vinifera* (cv. Shiraz, field grown) leaf with increasing doses (10 to 100 μL) of pure methanol added on a filter paper below the leaf. Gas exchange was simultaneously measured and VOC sensors indicate the VOC concentration over time at the leaf plane (VOC1) and 30 cm away (VOC2). Sap solution = WD-AS pH 6.0, PAR = 1290 μmol m^−2^ s^−1^. Repeat experiments (n=7) allowed the construction of a dose-response curve (b) where relative flow is plotted against the logarithm (base 10) of the parts per million (by volume, ppmv) of methanol measured at the leaf plain by VOC1 and integrated over time (ppmv × h). EC50 = 11.3 ppmv × h, R^2^ = 0.69. c) Dose response curve for methanol applied on the filter paper, EC50 = 43 μL R^2^ = 0.87.

After the transient detailed in Figure 8 there was a delay before the flow rate started to drastically decrease and then increase again until reaching a similar rate to the initial conditions prior to the addition (Figures 7, 9). This dramatic effect was also observed on leaves that displayed oscillating flow rates.

From several experiments with different methanol concentrations a dose response curve could be constructed for the *Q* response in *V. vinifera* leaves (Figure 9b, c). Dose responses could be constructed with either the amount added to the filter paper (Figure 9c) or the concentration detected at the level of the leaf surface (Figure 9b). Slightly better R^2^ were obtained with the amount added.

Arabidopsis leaves also responded to methanol but not ethanol similarly to *V. vinifera* (Figure 10). The response to 50 uL of methanol was on average a reduction to 40% (SD = 13%) of the initial flow before methanol was added.

**Figure 10.**
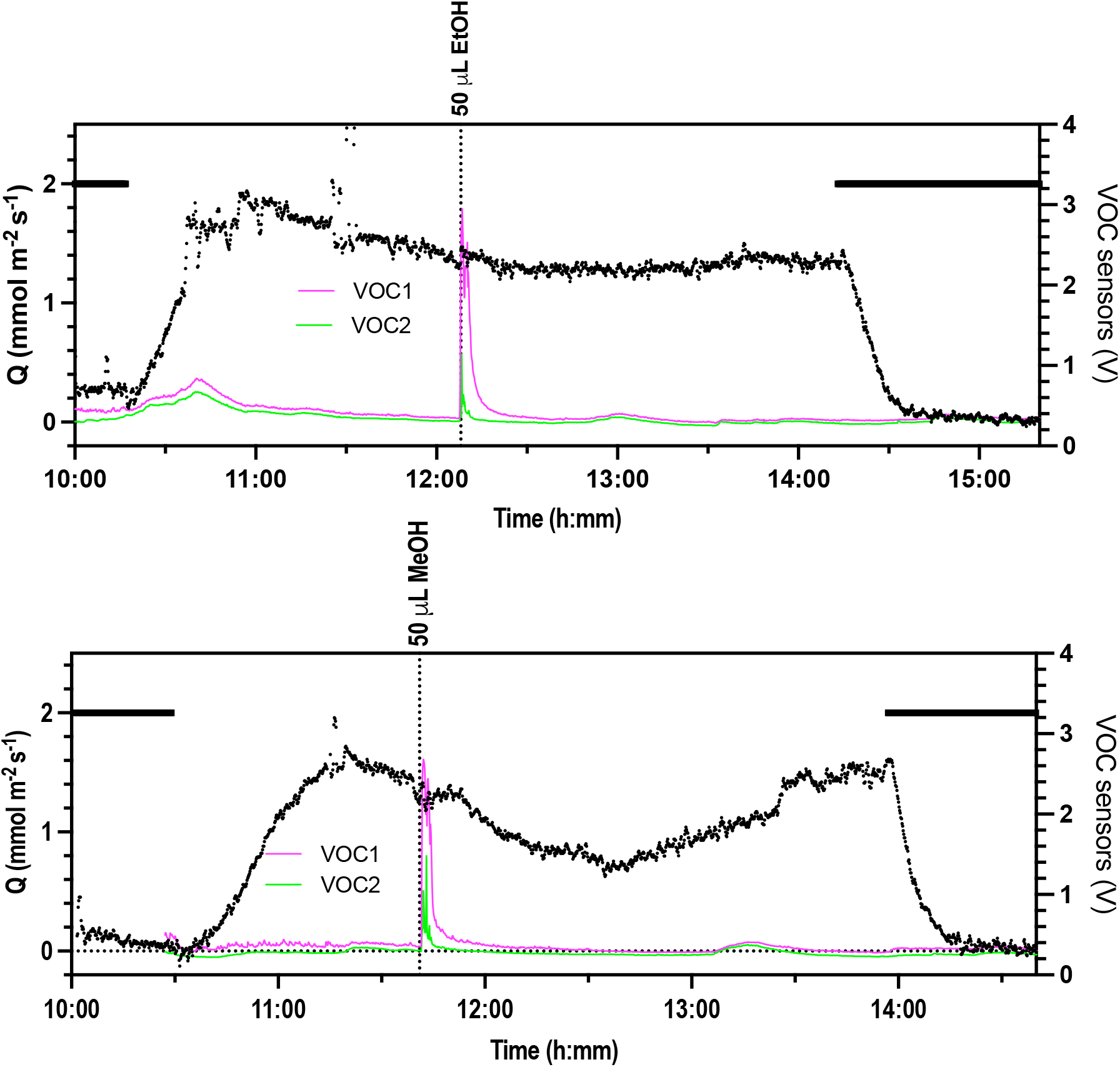
Response of flow (*Q*) into Arabidopsis leaves to ethanol and methanol applied as volatiles (VOC1 magenta, VOC2 green). Solid horizontal black line = dark period. Sap solution = WD-AS pH 4.5, PAR = 100 μmol m^−2^ s^−1^. Example traces from 3 repeats each.

### 3.7 Leaf-emitted volatiles during stomatal oscillations and dark transition

Repeated large oscillations were observed on a *V. vinifera* leaf connected to the flow meter, after 1.5 hour in one experiment (Supplementary Figure 5). As the gas sensors were placed under the leaf, the gas sensor 1 detected volatiles during the ascending phase of the oscillations while gas sensor 2 positioned 15 cm below the leaf did not. This likely indicates that VOCs were emitted from the leaf when stomata were open. Additionally, dark transitions also revealed a VOC release as gas sensor 1 (close to the leaf) detected volatiles but the gas sensor 2 did not (e.g. Figure 6b) and this was observed multiple times (n=3).

## 4. DISCUSSION

### 4.1 Methanol-induced stomatal closure

Our observations reveal two responses in transpiration to applied VOCs both of which are dependent on the VOC molecule and the dose. The first is a small transient reduction in *Q* that closely correlates with the transient in VOC concentration measured with the gas sensors close to the leaf surface (Figure 8). This can be attributed to the collision of the VOC molecules entering the leaf with water molecules leaving the leaf and is a factor also considered in estimating entry of other gases through stomata such as CO_2_ and ozone (Jarman, 1974; Uddling, Matyssek, Pettersson, & Wieser, 2012). In principle the response could be modelled given the characteristics of the VOCs applied, however we cannot assume that all the stomata on the leaf surface “see” the same VOC concentration and thus the response in *Q* is an average for the whole leaf. Nevertheless the initial response can be used to indicate that ethanol and methanol are entering the leaf in these experiments via the stomata; a conclusion also made by an earlier study on the effects of methanol fumigation on photosynthesis (Loreto, Tricoli, Centritto, Alvino, & Delfine, 1999).

The second slower response in *Q* to methanol that indicates that stomatal closure occurs after most of the methanol has diffused away from the leaf surface (e.g. Figures 7, 9, 10). This response does not occur with ethanol, or methyl salicylate that has been listed previously as a volatile that can cause stomatal closure (Agurla, Sunitha, & Raghavendra, 2020). No responses were observed to methanol in the isolated part of the leaf enclosed in the IRGA cuvette indicating that this is not a systemic response, at least in *V. vinifera*. The methanol response is dose dependent and indicates at least a partial stomatal closure then re-opening in both *V. vinifera* and Arabidopsis. It is unlikely to be due to a change in apoplastic pH since methanol does not change pH when in solution. Methanol is likely to be metabolised in the leaf and/or stomatal guard cells to produce formaldehyde, formate and CO_2_ (Fall & Benson, 1996) and these in turn may result in a stomatal closure response. On the other hand various signalling responses to volatile methanol have been reported in plants including an elevation in cytosolic calcium concentration and membrane depolarisation (Tran et al., 2018) that can trigger stomatal closure. A previous report examining the effect of methanol fumigation on some horticultural plants found no effect on stomatal conductance (Loreto et al., 1999), though another study found inhibited stomatal opening with methanol but at very high solution concentrations (Mouravieff, 1976).

Some volatiles are known to affect stomata, for example, gaseous hydrogen sulphide (H_2_S) was found to mediate stomatal movements (Du et al., 2019) and some green leaf volatiles were shown to induce the closure of stomata for pathogen defence responses in tomato (López-Gresa et al., 2018). These GLVs were tested here at the same concentrations by adding certain volumes on a filter paper close to the leaf on *V. vinifera* but showed no effect on flow rates. Volatile methanol shown here to cause a transient stomatal closure in Arabidopsis and *V. vinifera* is a highly water-soluble volatile that is well known to be emitted by plants (Jacob et al., 2005; Jud et al., 2018). Emission is triggered when there are changes in cell wall structures during seed maturation, fruit ripening, leaf expansion and biotic stress (Dorokhov, Sheshukova, & Komarova, 2018). Pectin methyl esterases are the likely instigators of methanol production (Dorokhov et al., 2018). A morning peak of methanol emission has been shown in normal conditions coinciding with an increase in stomatal conductance, followed by a slow and gradual decrease during the day (Hüve et al., 2007). Moreover, methanol has been found to induce defence reactions in intact leaves from the same and neighbouring plants, and to activate resistance genes (Dorokhov et al., 2012). Spraying leaves with methanol showed a stimulation of photosynthesis activity and productivity in C3 plants (Nonomura & Benson, 1992), and the regulation of genes involved in signalling, defence and metabolism in Arabidopsis (Downie et al., 2004). A rise in temperature also induced an increase in MeOH emission by up to 12% per degree, and a dark to light transition increased the MeOH emission by twofold (Folkers et al., 2008). Pathogen interactions induce the emission of MeOH, for example, during *Manduca sexta* caterpillar attack in *Nicotiana attenuate* (Von Dahl, Hävecker, Schlögl, & Baldwin, 2006), and during feeding of *Euphydryas aurinia* caterpillars on *Succisa pratensis* (Peñuelas, Filella, Stefanescu, & Llusià, 2005). Methyl jasmonate induces an increased emission of methanol from leaves of cucumber (Jiang, Ye, & Niinemets, 2021). However for barley methanol was considered to be mainly a constitutive emission rather than stress induced based on treatment with the elicitor benzothiadiazole (W. Jud et al., 2018). In summary methanol emission via stomata is common in plants linked to growth and stress (abiotic and biotic) but responses are likely to be species dependent. There appears to be no previous evidence of direct application of volatile methanol inducing a relatively rapid change in transpiration (stomatal conductance) as demonstrated here.

We are not aware of methanol being reported to be released by *V. vinifera* or Arabidopsis though for *V. vinifera* a large variety of other VOCs are released under drought stress (Griesser et al., 2015). Given its common occurrence as a released VOC from leaves it would be surprising if methanol were not released from *V. vinifera* and Arabidopsis. How this may relate to our observations of a dose dependent transient stomatal closure is not clear since methanol is released often as stomata open (Hüve et al., 2007). It may be that the dose applied in our case is much higher than the volatile concentrations that accumulate and are released from leaves, however even small concentrations in the vapour phase had a measurable closing effect and the methanol that enters via the stomata would have to enter the liquid phase to induce the closing effect. Alternatively the sensitivity to methanol may be dependent on the transition phase or state of the stomata, that is, stomata may be less sensitive to methanol during the opening phase when methanol is normally released. Other alternatives exist to be further explored including that methanol may be part of a feedback system or that stomatal responses may depend on a combination of methanol with other VOCs, for example, acetic acid (Jardine et al., 2022).

Gas sensors were added to the experimental system to monitor diffusion of volatiles during the experiment and to estimate when the leaf would detect the volatiles and when the volatiles would eventually disperse in the air. Nevertheless, the non-specific sensors used were also able to detect volatiles emitted by the leaf but not to the resolution of specific molecules as used in other studies e.g. (Jardine et al., 2022; Jud et al., 2018). They were detected from *V. vinifera* leaves during the light-dark transition but not dark-light as observed in other studies (Jud et al., 2016) though for barley light-dark transitions appear to show a small peak in methanol emission (Jud et al., 2018). The inexpensive gas sensor used here is more specific for alcohols, and could be calibrated against methanol, but it is also sensitive to other types of volatiles. During oscillations of transpiration rates measured by the flow meter we also observed VOC emissions on some occasions (Supplementary Figure 5). The gas sensors closest to the leaf detected volatiles during the peak of oscillations that then disappeared during the lower part of the oscillations or became too low in concentrations for the threshold sensitivity of the sensors. Thus, it strongly suggests that the volatiles were emitted by the leaf since the second gas sensor further from the leaf did not detect any. This observation could indicate a potential feedback role of volatiles in stomatal regulation. A volatile ion emitted from the leaf mesophyll that alters epidermal pH has been implicated in stomatal control (Mott et al., 2014) but its ionic nature and alteration of pH would exclude volatile methanol in this case.

### 4.2 Leaf water flow monitored with a flow meter

Quantifying stomatal responses is important to understand how plants respond to environmental stimuli. In this study, sensitive flow meters connected to petioles of single leaves of Arabidopsis and *V. vinifera* with suitable perfusate solutions provided a robust method for measuring changes in transpiration. The responses in flow (*Q*) were rapid to stimuli at the leaf level, e.g to light (Figures 2, 3, 4, 6), cavitation (Figures 5, 6), air flow (Figure 6) and VOCs (Figures 7, 8, 9, 10) and continuous measurements were conducted with steady flow rates that were recorded over at least a light period in most cases confirming that the cut of the petiole in the solution posed little effect when following the protocol described in Ceciliato et al. (2019). Oscillations in *Q* were also observed and are relatively common when monitoring stomatal conductance (Ballard, Peak, & Mott, 2019; Yang, Zhang, & Zhang, 2005).

Transpiration rates are generally determined from the water vapour released from open stomata on a leaf. However, the flow method described here provides measurements of the water entering the leaf through the petiole xylem vessels. In order to evaluate these flows as a measure of transpiration, a gas exchange system was simultaneously attached to the leaf. Combined with simultaneous flow measurement, at least for *V. vinifera*, it was possible to better resolve some signalling events within the leaf to external factors, since the part of the leaf under the IRGA cuvette is isolated from the treatments on the rest of the leaf and resolved by the flow meter. For example, the effect of methanol did not cause a change in *E* or *A* in the isolated part of the leaf measured by the IRGA (e.g. Figures 7, 9) indicating a local effect on only the part of the leaf exposed to methanol vapour.

Flow into the leaf may not necessarily correspond to leaf transpiration if the leaf is re-hydrating, growing or dehydrating. This was evident often after the light was turned off since *E* recorded with the IRGA would return to zero more rapidly than flow (*Q*). That is, the water volume of the leaf may not be constant in time during these experiments and therefore the flow rate may not correspond exactly to the transpiration rate. Rarely was there an exact match between flow and transpiration for *V. vinifera* leaves noting that it is well known that stomatal conductance is patchy in *V. vinifera* (Düring & Stoll, 1996) and combined with only a relatively small fraction of the total leaf area incorporated in the IRGA cuvette, this could easily account for a mismatch. Compared to the amount of water in a *V. vinifera* leaf (average 2.44 +/− 0.72 (SD) g, n=24) typical steady flow rates that were measured (2.93 g h^−1^ +/− 0.74 (SD) g h^−1^, n=24) would mean that the leaf water was turned over on average every 53 min (range 29 to 184 min). For Arabidopsis leaves a match was more common since most of the leaf area was enclosed in the IRGA cuvette. This comparison is confounded by the effect of applying the IRGA cuvette to the leaf where the seal pressure on the leaf surface may cause a disruption in vascular continuity. For Arabidopsis as soon as the LCpro-SD cuvette was placed on the leaf, readings from the flow meter were seriously disrupted, probably because the xylem conduits were being squeezed by the cuvette seals. It should be noted that if only the IRGA were being used this effect would not be seen by the experimenter (e.g. Figure 3, Supplementary Figure 2).

### 4.3 Generation of xylem tension within *V. vinifera* leaves and the response to hydraulic failure

By incorporating a hydraulic resistance in series with the petiole connection it was possible to deduce the pressure at the entry of flow to the cut end of the petiole. At high flow rates induced by leaf transpiration these pressures were negative and therefore closer to a physiologically relevant state. Considering the remainder of the petiole hydraulic conductance measured at the end of the experiment we could deduce the pressure at the junction of the petiole and leaf lamina in *V. vinifera*. These experiments were not attempted for Arabidopsis since flow rates were two low to induce negative pressures with the hydraulic resistance available.

The deduced leaf xylem pressures (*LXP* in the Figures 2, 5, 6, 7, 8) were often negative (~ −1.0 bar), depending on supply pressure and flow rate. Gradually lowering the supply pressure or inducing greater flow with increased airflow over the leaf surface would often cause hydraulic failure presumed to be a cavitation somewhere in the system, which is indicated by a sudden reduction in *Q* and onset of leaf wilting. This always resulted in an immediate increase in *E* and *A* recorded with the IRGA and presumed to be caused by a sudden increase in cell turgor as xylem tension in the leaf is suddenly reduced. As water is lost, turgor then declines and both *E* and *A* decline rapidly. This can be reversed by increasing the supply pressure that initially results in relatively high flow close to that of the initial rate and then declines transiently at constant supply pressure as *E* and *A* show a transient recovery. During this phase the apparent leaf xylem pressure is calculated to be negative, however this is unlikely to be the case. The hydraulic failure (cavitation) is most likely to have occurred within the PEEK tubing and probably towards the petiole junction where tensions would be the most negative. The hydraulic resistance of the PEEK tubing used in the calculation is only valid if the tubing is completely full of fluid. When a cavitation occurs in the tube and/or petiole, the resulting vacuum would expand releasing sap to the leaf but the actual hydraulic conductance of the tubing would be higher than when it is full of sap. Thus it is likely that the tensions calculated when refilling occurs are overestimated. The point where leaf xylem pressure reaches a maximum and then declines with recovery of *E* and *A* is more likely to be a true reflection of the actual leaf xylem pressure. The continued flow measured after a presumed cavitation and during leaf wilting is due to the pressure gradient of a vacuum (or close to depending on vapour pressure) in the cavitation and the applied pressure in the instrument.

Another important observation from these experiments to be further explored was the relative constancy of flow and transpiration rate despite variation in xylem pressure and supply pressure. This is most clearly seen in the experiments shown in Figure 5, but was observed on multiple occasions with a wide variation in supply pressure. Flow also returned close to the initial value after the expected transients in response to a change in air flow across the leaf (Figure 6). This response is expected to be complex since feedback systems change stomatal conductance in response to altered CO_2_ concentration and water vapour concentration resulting from changes in boundary layer conductance (Aphalo & Jarvis, 1993).

### 4.4 Conclusion and prospects

We have demonstrated that volatile methanol, but not ethanol or methyl salicylate (*V. vinifera*) can induce a localised dose dependent closure of stomata after entering the leaf via the stomata. It would be interesting to survey a range of plant species to determine the generality or otherwise of the methanol response. That Arabidopsis shows this response allows interrogation of the various stomatal closing mutants that are available that may reveal the signalling pathway involved. The method described here for monitoring flow at high sensitivity and with negative xylem tensions akin to the intact state of a leaf was advantageous for monitoring the responses in transpiration to external factors such as light, air movement and volatiles. In addition, the method allowed to investigate how leaves are actually responding to being connected to IRGAs and demonstrating for Arabidopsis a drastic reduction and erratic flow rate measured with the flow meter despite apparent normal behaviour in IRGA-measured *E* and *A*. Responses to xylem pressure and cavitation under the control of the experimental system can also be readily monitored in real time and future studies combining sensors for leaf xylem emboli (Brodribb et al., 2016) may reveal important responses in stomatal conductance and leaf hydraulic conductance.

## Acknowledgements

We acknowledge the assistance of staff in the Adelaide Plant Accelerator, Dr Sunita Ramesh for assistance with Arabidopsis plants. The research was funded by Australian Research Council Centre of Excellence in Plant Energy Biology CE140100008.

## Data availability statement

The raw data from all experiments as well as the material used in this manuscript can be obtained from the corresponding author upon reasonable request.

## Supplementary information

Supplementary Figure 1. Calibration of VOC sensors.

Supplementary Figure 2. Effect of attachment of the IRGA cuvette onto Arabidopsis leaves.

Supplementary Figure 3. Light-dark-light transistions measured on glasshouse gown *Vitis vinifera*.

Supplementary Figure 4. Induction of hydraulic failure (cavitation) and wilting by increasing airflow.

Supplementary Figure 5. Flow rate (Q) measured for a *V. vinifera* leaf (cv. Shiraz, glasshouse grown) connected to the flow meter where large oscillations were recorded corresponding to oscillations in volatile emissions from the leaf.

Supplementary S2. Arduino gas sensors program code

